# Monoclonal anti-dsRNA antibody-based metagenomics (MADAM) reveal *Pyricularia oryzae* mycovirome

**DOI:** 10.64898/2026.05.18.725940

**Authors:** Laurence Blondin, Denis Filloux, Emmanuel Fernandez, Henri Adreit, Huichuan Huang, Elisabeth Fournier, Didier Tharreau, Philippe Roumagnac

## Abstract

This study introduces MADAM (Monoclonal Anti-dsRNA Antibody-Based Metagenomics), a novel approach that integrates multiple technical modules previously used independently in other protocols. MADAM combines monoclonal antibody-mediated double-stranded RNA (dsRNA) enrichment, sequence-independent RT-PCR, and Oxford Nanopore Technologies (ONT) sequencing. Applied to *Pyricularia oryzae*, the causal agent of rice blast disease, MADAM enabled the comprehensive characterization of mycovirus genomes from four fungal isolates collected in Yunnan, China. The approach achieved high viral read recovery rates (46.9-72.7%) and identified 18 P*. oryzae*-associated RNA viruses spanning seven families: *Botourmiaviridae*, *Deltaormycoviridae*, *Mymonaviridae*, *Partitiviridae*, *Polymycoviridae*, *Splipalmiviridae*, and *Ambiguiviridae*. Seventeen nearly complete to complete viral genomes (1,226-6,085 nucleotides) were recovered, with sequence coverage ranging from 88% to 100%. Co-infections were detected in three of the four isolates, with notable discoveries including the first deltaormycovirus reported in *P. oryzae*, a putative novel member of *Botourmiaviridae*, and an additional genomic segment of a polymycovirus. MADAM successfully detected positive-sense, negative-sense ssRNA, and dsRNA viruses, demonstrating its broad applicability. By uncovering novel viruses and resolving complex co-infections, this method proves invaluable for fungal virology, with potential applications in diagnostics, surveillance, and biological control. Ultimately, MADAM advances our understanding of fungal viral diversity and paves the way for further exploration of mycovirus ecology and evolution.

## Introduction

Mycoviruses, which infect and replicate within fungal cells, were first identified in *Agaricus bisporus* in 1962 (Hollings 1962). Initially studied primarily in ascomycetes and basidiomycetes, they have since been detected in early-diverging fungal lineages, expanding our understanding of their ecological and evolutionary significance (Xie & Jiang 2014; Myers *et al*. 2020). These viruses have garnered increasing interest due to their potential as biological control agents. Some mycoviruses induce hypovirulence, reducing the pathogenicity of phytopathogenic fungi and offering sustainable alternatives to chemical fungicides (Wagemans *et al*. 2022; Galli *et al*. 2024). Others trigger hypervirulence, enhancing fungal antagonism against insect pests or weeds (Guo *et al*. 2024). Additionally, the discovery of hypovirulence-associated mycoviruses in murine and human-pathogenic fungi presents novel opportunities for antifungal therapies (Takahashi-Nakaguchi *et al*. 2020; Battersby *et al*. 2024).

The International Committee on Taxonomy of Viruses (ICTV) has officially classified over 674 mycovirus species spanning over 53 families and 104 genera (VMR MSL41.v1, March 20, 2026; available https://ictv.global/vmr), but many newly identified mycoviruses remain unassigned. The vast majority of characterized mycoviruses possess genomes composed of either double-stranded RNA (dsRNA) or positive-sense single-stranded RNA ((+)ssRNA). In contrast, negative-sense single-stranded RNA ((−)ssRNA) and DNA viruses represent only a minor fraction of the known mycovirus diversity (Xie & Jiang 2024). Single-stranded DNA (ssDNA) mycoviruses, in particular, are still understudied, likely due to their low abundance or methodological detection biases (Myers *et al*. 2020). Furthermore, co-infections, where multiple viruses infect a single host, appear to be the rule rather than the exception, giving rise to complex and unpredictable interactions between viruses and between viruses and their fungal hosts (Thapa & Roossinck 2019; Ayllón & Vainio 2023). For example, a recent large-scale study of *Botrytis cinerea* revealed that 404 out of 406 field isolates harbored between 2 and 25 distinct mycoviruses (Duan *et al*. 2025), highlighting the ubiquity and complexity of mixed viral infections in natural fungal populations.

Addressing the challenges of mycovirus diversity and community composition analysis has driven substantial methodological advancements in their detection and characterization. These methods include the virion-associated nucleic acid (VANA) approach, small RNA sequencing, metatranscriptomic approaches and double-stranded RNA (dsRNA)-based techniques (Roossinck *et al*. 2015). Among these, dsRNA sequencing stands out due to its ability to selectively enrich viral genomes and replication intermediates, thereby reducing host RNA/DNA background noise and improving sensitivity and genome coverage (Decker *et al*. 2019; Gaafar & Ziebell 2020; Javaran *et al*. 2023).

The classic dsRNA extraction method, based on cellulose chromatography (Morris & Dodds 1979), has been widely used for over five decades to isolate viral dsRNA from infected plant, animal, and fungal samples. While this approach has been refined to improve yield and purity (Marais *et al*. 2018a; Cardoso *et al*. 2023; Wang *et al*. 2024), it remains labor-intensive, limiting its scalability. Despite this, it has successfully enabled the reconstruction of full viral genomes in key phytopathogenic fungi and oomycetes, such as *Cryphonectria parasitica* (Hillman & Suzuki 2004), *Botrytis cinerea* (Drury *et al*. 2025), *Fusarium* spp. (Buivydaitė *et al*. 2024), *Rosellinia necatrix* (Hassan *et al*. 2025), *Sclerotinia sclerotiorum* (Contreras-Soto & Tovar-Pedraza 2023) and *Phytophthora* spp. (Cai & Hillman 2013).

Despite recent dsRNA-based technique improvements, such as the novel cellulose-magnetic bead-based method using ReliaPrepTM Resin (Rott *et al*. 2024), novel enrichment strategies have emerged. One approach uses the dsRNA-binding protein B2 from Flock House virus, which simplifies purification and enhances recovery efficiency (Rott *et al*. 2024; Fall *et al*. 2025). An alternative approach leverages anti-dsRNA monoclonal antibodies (mAbs), which exhibit exquisite specificity for dsRNA, including the ability to detect fragments as short as ∼14 base pairs (Bou-Nader *et al*. 2025). Detailed biophysical characterization has elucidated the molecular basis of this interaction, demonstrating that J2 mAb achieves this targeted interaction by deploying both its heavy and light chain complementarity-determining regions in a coordinated manner, effectively tracking the minor groove of the dsRNA helix (Bou-Nader *et al*. 2025). This dual-chain engagement underpins the antibody’s precise and selective recognition of dsRNA molecules (Bou-Nader *et al*. 2025). Antibodies were originally developed to detect animal viruses (Richardson *et al*. 2010; O’Brien *et al*. 2015; Decker *et al*. 2019) but have since been adapted for plant viruses (Blouin *et al*. 2016; Blouin *et al*. 2023). For example, the 2G4 mAb-based protocol successfully identified known plant viruses (e.g., potato virus Y, turnip mosaic virus) and even discovered a novel macluravirus (Blouin *et al*. 2016).

In this study, we used a monoclonal anti-dsRNA antibody-based metagenomics (MADAM) protocol that integrates dsRNA enrichment via the 2G4 monoclonal antibody, sequence-independent RT-PCR amplification of viral cDNA, and Oxford Nanopore Technologies (ONT) to enable rapid and comprehensive characterization of mycovirus genomes in *Pyricularia oryzae* (syn. *Magnaporthe oryzae*), the fungal pathogen responsible for rice blast disease. This approach enabled the detection of previously characterized viruses with high genome coverage, while also revealing novel viral entities, including a putative novel ssRNA virus (*Botourmiaviridae*), an additional genomic segment of a multi-segmented polymycovirus, and the first deltaormycovirus reported in *P. oryzae*. To ensure complete genome assembly of both known and novel viruses, 5ʹ and 3ʹ terminal sequences using a modified 3ʹ RACE approach were further obtained.

## Materials and Methods

### Origin, growth conditions, and harvest of the fungal isolates

Four *Pyricularia oryzae* strains, isolated from rice lesions (panicle or leaf) collected from the Yuanyang rice terrasses (Yunnan, China) were used in this study (Supplementary Table 1). Conidia from 10-day-old cultures were harvested, counted using a Malassez chamber and 1 x 10^6^ spores were inoculated into 200 ml of liquid TNK-YE-Glucose medium (10g/l glucose, 2 g/l yeast extract, 2 g/l NaNO_3_, 2 g/l KH_2_PO_4_, 0.5 g/l MgSO_4_-7H_2_O and 0.1 g/l CaCl_2_- 2H_2_O, 0.004 g/l FeSO_4_7H_2_O and 1 ml/l of a stock solution of microelements (7.9 g/l ZnSO_4_-7H_2_O, 0.6 g/l CuSO_4_-5H_2_O, 0.1 g/l H_3_BO_3_, 0.2 g/l MnSO_4_-H_2_O and 0.14 g/l NaMoO_4_-2H_2_O), pH was adjusted to 5.5–5.8) and incubated at 110 rpm, 25°C for 48h. The resulting mycelia were filtered through Miracloth, digested with ß-glucanase (Extralyse-Laffort) for 2h and the released protoplasts obtained were collected by centrifugation (2,000 rpm, 10 min). The supernatant was discarded and the protoplast pellets were lyophilized for downstream analysis.

### Virion-associated nucleic acid (VANA) metagenomics

Samples from two strains (CH2054 and CH2061) were processed using the virion-associated nucleic acid (VANA) metagenomic approach (François *et al*. 2018), using mycelium obtained as described above. Sequencing was performed using an Illumina HiSeq 3000 platform by Azenta (South Plainfield, NJ, USA). The bioinformatic analyses were conducted as previously described (Moubset *et al*. 2024).

### Monoclonal anti-dsRNA antibody metagenomics (MADAM)

Lyophilized protoplasts were ground into a fine powder using a 1600-MiniG homogenizer (1,500 rpm, 2 min). Total RNA was extracted following the protocol described by Ballini et al. (Ballini *et al*. 2013) with minor modifications. After phenol extraction and centrifugation, chloroform–isoamyl alcohol was added to the aqueous phase. RNA was subsequently precipitated with isopropanol in the presence of sodium acetate (final concentration 0.3 M). Nucleic acids (DNA and single-stranded RNA) were removed by nuclease treatment using RNase T1 (Thermo Fisher Scientific) and DNase I Ambion (Invitrogen). Then, double-stranded RNA was enriched using the monoclonal antibody anti-dsRNA- mAb 2G4 (UniQuest Pty Limited) as described by Blouin et al. (Blouin *et al*. 2016).

Strand-switching cDNA synthesis from dsRNA templates was performed using a modified ONT PCR-cDNA protocol (SQK-PCS109). Random octamer primers (VNP8N, 20 µM; Eurogentec) containing ONT anchor sequences (Supplementary Table 2) replaced the polyT primer. Briefly, 10 µL of resuspended dsRNA-beads was denatured at 95°C for 2 min, chilled on ice for 2 min, and subjected to reverse transcription using a reaction mix containing 200 U Maxima H Minus Reverse Transcriptase (Thermo Fisher Scientific), 0.5 mM dNTPs, 4 µL of 5X RT buffer, 40 U RNaseOUT (Invitrogen), and 2µl of 10 µM strand-switching oligo (SSP, Supplementary Table 2). The reaction proceeded at 25°C for 10 min, followed by 42°C for 1.5 h and 85°C for 5 min. Ribonuclease treatment (1 µL RNase Cocktail, Invitrogen) was performed at 37°C for 10 min, and cDNA was purified using AMPure XP beads (Beckman Coulter) at 0.8X ratio, and eluted in 20 µL High-Pure water.

For second-strand synthesis, two parallel reactions were prepared: 10 µL of purified cDNA was combined with 1 µL VNP8N primer (20 µM), and another 10 µL was combined with 1 µL PR2 primer (10 µM; complementary to SSP, Supplementary Table 2). Both reactions were denatured at 95°C for 2 min and rapidly cooled to 4°C for 2 min. Thus, 0.5 µL Klenow DNA polymerase (New England Biolabs), 1× Klenow reaction buffer (NEB2), and 0.7 mM dNTPs were added, followed by incubation at 16°C for 90 min and enzyme inactivation at 75°C for 10 min. The reactions were pooled, and branched DNA intermediates were resolved using 1 µL T7 endonuclease (New England Biolabs) at 37°C for 30 min. The final double-stranded cDNA was purified using AMPure XP beads (Beckman Coulter) at 0.8X ratio and eluted in 10 µL High-Pure water.

PCR amplification was performed individually using 5 µL of Klenow-synthesized cDNA in two parallel 25 µL reactions with LongAmp Taq DNA Polymerase (New England Biolabs). Each reaction contained 12.5 µL of 2X LongAmp buffer, 1 µL of 3580R primer (10 µM), and 1 µL of PR2 primer (10 µM) (Supplementary Table 2), which are complementary to the VNP8N and SSP primer anchors, respectively. Thermal cycling conditions included an initial denaturation at 95°C for 30 s, followed by 35 cycles of 95°C for 30 s, 62°C for 30 s, and 65°C for 3 min, with a final extension at 65°C for 6 min. The amplified products from both reactions were pooled and analyzed on a 1% agarose gel to evaluate amplification efficiency. Finally, 1 µL of Exonuclease I (New England Biolabs) was added to degrade residual primers prior to bead purification at 0.7X ratio (AMPure XP ratio, Beckman Coulter).

MinION libraries were prepared from each amplicon using the SQK-LSK114 kit (Oxford Nanopore Technologies) according to the manufacturer’s protocol. Briefly, DNA underwent end repair and dA-tailing using the NEBNext End Repair/dA-Tailing Module (New England Biolabs), followed by bead purification (1.0× bead-to-volume ratio). Sequencing adapters (Oxford Nanopore Technologies) were ligated to end-prepared DNA using NEBNext Quick T4 DNA Ligase (New England Biolabs), and libraries were purified using AMPure XP beads (0.4× ratio) with ONT Short Fragment Buffer. Each library (∼30 fmol) was combined with ONT sequencing buffer and loading beads, then loaded onto FLO-FLG114 R10.4.1 Flongle flow cells (Oxford Nanopore Technologies). Sequencing of each library was performed separately in simplex mode on a MinION Mk1B device using MinKNOW v25.09.16 with a 24-hour run protocol.

### Completion of genomic sequences

Target cDNAs were synthesized and amplified from dsRNA fragments using gene-specific primers (Supplementary Table 2) with the One-Step RT-PCR Kit (Qiagen), following the manufacturer’s instructions. Amplicons were either Sanger-sequenced (Genewiz) or purified, pooled, and sequenced on a Flongle flow cell using MinION library preparation, as described below.

To resolve 3ʹ and 5ʹ termini of dsRNA molecules when random-primed cDNA synthesis yielded incomplete coverage, a similar Nanopore cDNA-PCR (SQK-PCS109) strategy was employed. Firstly, dsRNA was denatured at 95°C for 3 min, chilled on ice for 2 min, and polyadenylated at 37°C for 45 min using Poly(A) Polymerase (New England Biolabs), followed by AMPure bead purification (1× ratio). Then, first-strand cDNA synthesis was performed using Maxima H minus (Thermo Fisher Scientific) at 42°C for 90 min (inactivated at 85°C for 5 min) with a custom primer (0.1 µM; Eurogentec), analogous to the Nanopore VNP primer (15-poly(T) tail with a 5ʹ Nanopore anchor and 3ʹ terminal ‘V’), alongside the strand-switching oligo (SSP; Supplementary Table 2). Amplicons were generated via PCR amplification (PR2 and 3580R primers) as previously described, followed by Nanopore Flongle sequencing.

For unresolved cases, an alternative method (Brown *et al*. 2013) was applied to determine terminal sequences of dsRNA genomes and ssRNA replicative intermediates. cDNA-poly(A) extremities generated as described above, excepted using SuperScriptIII (Invitrogen) at 42°C for 60 min (inactivated at 70°C for 5 min), were PCR-amplified using 0.4 µM of a VNP-anchor complementary primer (3580R) and 0.4 µM of virus-specific primers (Supplementary Table 2), targeting 300–700 bp fragments. Amplification was performed with LongAmp Taq (New England Biolabs) under the following conditions: 95°C (30 s); 35 cycles of 94°C (30 s), 62°C (30 s), and 65°C (1 min); and a final extension at 65°C (10 min). Purified PCR products (AMPure beads, 1× ratio) were pooled and sequenced on Flongle flow cells using standard MinION library preparation.

### Bioinformatics analyses

The bioinformatics analysis of the Nanopore reads was carried out using the in-house pipeline MinIONMaster (available online at https://zenodo.org/records/20205917), which consists of the following steps: super-accurate basecalling was initially performed using Dorado v1.3.0 (available at: https://github.com/nanoporetech/dorado/releases/tag/v1.3.0). This was followed by adapter and primer trimming, as well as the removal of reads shorter than 50 nucleotides and reads with a Phred quality score below 7, using MinIONCleaner, a module of the MinIONMaster pipeline. Finally, taxonomic assignment was achieved on cleaned Nanopore reads through searches against the NCBI nr protein database using DIAMOND v2.1.6 (Buchfink et al. 2015) with an E-value threshold of < 0.001.

### Genome annotation of novel viruses

De novo assembly of read clusters, each potentially representing distinct viral families or genera, was performed in Geneious v6.1.8 using the De novo Assembly function with medium sensitivity (≤5 iterations, 75% similarity threshold and in some case 50% strict similarity threshold) to generate consensus sequences. These consensus sequences were then subjected to BLASTn and BLASTx searches against the NCBI nr database under default parameters. The top hit, defined by the highest query and subject coverage, was selected as the reference for each cluster. Reads were realigned to the reference, and consensus sequences were iteratively refined. Mean read depth and genome coverage were calculated using Geneious’ built-in statistics tool. Putative viral open reading frames (ORFs) were identified using NCBI ORF Finder (https://www.ncbi.nlm.nih.gov/orffinder/).

### Phylogenetic analyses

Amino acid sequences of complete RNA-dependent RNA polymerases (RdRps) from known and newly identified mycoviruses were aligned using MAFFT v7 (Katoh & Standley 2013). The alignment included reference RdRp sequences from ICTV-classified viral families and top BLASTp hits retrieved from the NCBI database, incorporating up to three representative sequences per genus, where possible. Maximum-likelihood phylogenetic trees were inferred using the NGS Phylogeny.fr platform (Lemoine *et al*. 2019) with the PHYML-SMS algorithm. Node robustness was evaluated using 1000 bootstrap replicates. Branches with less than 40% bootstrap support have been collapsed with TreeGraph2 (Stover & Muller 2010). Trees were visualized and annotated using iTOL (Letunic & Bork 2016).

## Results

### VANA metagenomics

The RNA and DNA virome of two *P. oryzae* isolates was characterized using the VANA approach. This metagenomic analysis identified one and eight partial RNA mycovirus genomes in the CH2054 and CH2061 *P. oryzae* strains, respectively (Supplementary Table 3). Notably no DNA mycoviruses were detected using this method. All nine mycoviruses were additionally confirmed using the MADAM approach and are fully described below.

### Eighteen *P. oryzae* associated RNA viruses revealed by the MADAM approach

The RNA virome of four *P. oryzae* isolates was characterized using the MADAM approach combining anti-dsRNA monoclonal antibody-based enrichment and Nanopore sequencing. Each isolate yielded 642,776 to 800,203 filtered reads, of which 46.9% to 72.7% were of viral origin (Table 1). Genome gaps were resolved for selected viruses via RT-PCR (using sequence-specific primers) and Sanger/Nanopore sequencing. However, Nanopore sequencing may not consistently capture the 5ʹ and 3ʹ terminal regions, which are critical for viral replication and propagation. To address this limitation, the terminal sequences were determined or validated using a modified 3ʹ RACE-PCR approach (Supplementary Table 2).

**Table 1:** Nanopore viral read count and sequencing depth mapped to taxonomically assigned contigs.

The complete sequences obtained (ranging from 1,226 to 6,085 nucleotides) achieved 88%–100% genome coverage relative to reference GenBank entries, except for a putative novel ssRNA virus (*Botourmiaviridae*) and an additional genomic segment of a multi-segmented polymycovirus. Co- infections were detected in three of the four *P. oryzae* isolates, each hosting at least three distinct mycoviruses (Table 1 and Supplementary Table 4).

BLASTn and BLASTp analyses revealed that all detected viruses shared significant sequence identities with previously described mycoviruses belonging to the families *Botourmiaviridae*, *Deltaormycoviridae*, *Mymonaviridae*, *Partitiviridae*, *Polymycoviridae*, *Splipalmiviridae*, and the *Ambiguiviridae* family, that was recently created by the ICTV to accommodate viruses previously known as mycotombusviruses (Yang *et al*. 2024).

For the remainder of the manuscript, we will use the prefix Po (for *Pyricularia oryzae*) as the host name in the nomenclature of the newly identified viruses. However, we will also frequently refer to the corresponding reference names registered in GenBank, which were originally assigned using the former host name, *Magnaporthe oryzae* = Mo prefix.

### Positive-sense single-stranded RNA viruses

#### Botourmiaviruses

Three and five botourmiaviral genomes were identified in the *P. oryzae* isolates CH1889 (named Pyricularia oryzae botourmiavirus 2 (PoBV2), PoBV3 and PoBV6-1; Table 1 and Supplementary Table 4) and CH2061 (PoBV4, PoBV6-2, PoBV7, PoBV14 and PoBV16; Table 1 and Supplementary Table 4), respectively. The high amino acid sequence identities observed in the RdRp regions of six of these genomes fell below the established species demarcation threshold (<90% RdRp amino acid identity), thereby supporting their classification within the same species as their respective closest reference viruses (Table 2). In addition, PoBV14 RdRp respectively demonstrated 98% and 97.6% aa identity with Xiaogan botourmia-like virus 2 (accession UUW21043) and Magnaporthe oryzae botourmiavirus 14 (accession XNZ83248), a virus recently described as a new member of the genus *Gammascleroulivirus* within the family *Botourmiaviridae* (Li *et al*. 2025). However, PoBV14 RdRp showed only 53% with the closest recognized reference virus Magnaporthe oryzae botourmiavirus 6 (accession QVU39993, species *Gammascleroulivirus alphaoryzae*) (Table2). Finally, PoBV16 RdRp shared 53.7% aa identity with Bremia lactucae-associated ourmia-like virus 2 (accession QIP68026) and 54.2% aa identity with Verticillium dahliae magoulivirus 1 (accession UVD54632) (Table 2), both botourmiaviruses being still unassigned to a specific genus by the ICTV. The 2,502-nucleotide genome of PoBV16 encodes a 623- amino-acid RdRp containing canonical viral polymerase motifs (Supplementary Figure 1). Phylogenetic analysis based on RdRp sequences of the 8 botourmiaviruses here described and representative members of all genera of the *Botourmiaviridae* family (Figure 1A) further suggest that both PoBV14 and PoBV16 may represent members of two putative novel genera of the *Botourmiaviridae* family.

**Figure 1.** **(A)** A maximum-likelihood phylogenetic tree was constructed based on the RdRp amino acid sequences of the *Botourmiaviridae* family genomes identified in this study (highlighted in red) and representative members of the *Botourmiaviridae* family available in GenBank. The analysis includes ICTV-validated genomes from all 9 fungal-infecting genera within the family. Node support was assessed using 1000 bootstrap replicates. **(B)** A maximum-likelihood phylogenetic tree was constructed based on the RdRp amino acid sequences of the *Bormycovirales* order including *Deltaormycoviridae* family genomes identified in this study (highlighted in red) and representative members of the *Bormycovirales* order available in GenBank. The analysis includes ICTV-validated genomes of different genera within the families. Node support was assessed using 1000 bootstrap replicates.

**Table 2.** RdRP amino acid sequence of the mycoviruses and their sequence similarities to closest available ICTV-recognized reference viral sequence or best-NCBI GenBank matching viral sequence.

#### Deltaormycoviruses

Two bipartite single-stranded RNA viral genomes were recovered from *P. oryzae* isolates CH1889 (named Pyricularia oryzae deltaormycovirus 1-1, PoDOmV1-1) and CH2061 (PoDOmV1-2), respectively (Table 1 -Supplementary Table 4 and Supplementary Figure 2A). RNA1 segments of PoDOmV1-1 and PoDOmV1-2 shared 69.6% and 69.4% (coverage 96%) nucleotide identity with Verticillium dahliae ormycovirus 1 (accession OR734290, species *Bormycovirus verticilli*), respectively, while RNA2 segments exhibited 75.7% and 75.4% (coverage 88-89%) nucleotide identity with the corresponding segments of Verticillium dahliae ormycovirus 1 (accession OR734291). Notably, the 5′and 3’ untranslated regions (UTRs) of both segments were highly conserved (Supplementary Figure 2B-C). At the amino acid level, RNA1-encoded RdRps of PoDOmV1-1 and PoDOmV1-2 shared 65.7% and 65.3% (coverage 100%) identity with the RdRp of Verticillium dahliae ormycovirus 1 (accession WPV08068) (Table 2), respectively, while their RNA2-encoded hypothetical proteins (HPs) displayed 74.4% and 74.6% (coverage 96%) identity with the corresponding HP of Verticillium dahliae ormycovirus 1 (accession WPV08069). The RdRp of PoDOmV1-1 and PoDOmV1-2. were 99.2% identical to each other. Consistent with the ormycovirus hallmarks, their *RdRp* gene contained the conserved ’NDD’ motif at position 486–488 (Supplementary Figure 2D). Phylogenetic analysis based on RdRp amino acid sequences (Figure 1B) revealed that PoDOmV1-1 and PoDOmV1-2 clustered with members of the *Bormycovirus* genus. Given the species demarcation threshold (<90% RdRp amino acid identity) for this genus, we propose that PoDOmV1-1 and PoDOmV1-2 represent isolates of a novel species of the *Bormycovirus* genus.

#### Splipalmiviruses

A multi-segmented viral genome (named Pyricularia oryzae splipalmivirus 1, PoSpV1), comprising four genomic segments, was identified in the *P. oryzae* isolate CH1889 (Table 1 and Supplementary Table 4). These segments shared 96.4–98.2% nucleotide identities with the corresponding segments of Magnaporthe oryzae narnavirus 1 strain J-YC (RNA1: accession LC553711, RNA2: accession LC553710, RNA3: accession LC553712 and RNA4: accession LC553713) (Table 1). belonging to the species *Delepalmivirus magnaporthae*. It is noteworthy that, in 2024, splipalmiviruses, phylogenetically related to narnaviruses but distinguished by split RdRPs encoded across two distinct genomic segments, were classified under the newly established family Splipalmiviridae. Consequently, Magnaporthe oryzae narnavirus 1 and Pyricularia oryzae splipalmivirus 1 are now regarded as synonymous virus names. Consistent with other *Delepalmivirus* members, PoSpV1 encoded a split RdRp domains distributed across two independent ORFs, including a 795 aa RdRp on RNA1 and a 783 aa RdRp on RNA2. The remaining segments, RNA3 and RNA4, encoded hypothetical proteins of 607 aa and 350 aa, respectively (Supplementary Figure 3A). Comparative alignment with splipalmiviruses confirmed the conserved division of RdRp domains, with motifs A, and B on one segment and motifs C and D on the other (Supplementary Figure 3C). Additionally, the RNA3 and RNA4 proteins of PoSpV1 exhibited 98.4% and 98.2% aa identities with their Magnaporthe oryzae narnavirus 1 counterparts (accessions BCH36657 and BCH36658), respectively. The 5′ and 3’ ends of all four segments were highly conserved (Supplementary Figure 3B), supporting their origin from a single viral entity. The high RdRp amino acid sequence identities observed for RNA1 and RNA2 genomes met the established species demarcation thresholds (<70% amino acid identity for both subunits), supporting their assignment to the same species as their respective closest reference viruses, *Delepalmivirus magnaporthae* (Table 2). Consistenly, phylogenetic analysis based on the RNA1 and RNA2 amino acid sequences (Figure 2A-B) reveal that PoSpV1 clustered closely with *Delepalmivirus magnaporthae*, tentatively indicating it represents an isolate of this species.

**Figure 2.** **(A)** A maximum-likelihood phylogenetic tree was constructed based on the RdRp amino acid sequences of the *Splipalmiridae* family genomes identified in this study (highlighted in red) and representative members of the *Splipalmiridae* family available in GenBank. The analysis includes ICTV- validated genomes of different genera within the family. Node support was assessed using 1000 bootstrap replicates. **(B)** A maximum-likelihood phylogenetic tree was constructed based on the RNA2 amino acid sequences of the *Splipalmiridae* family genomes identified in this study (highlighted in red) and representative members of the *Splipalmiridae* family available in GenBank. The analysis includes ICTV-validated genomes of different genera within the family. Node support was assessed using 1000 bootstrap replicates. **(C)** A maximum-likelihood phylogenetic tree was constructed based on the RdRp amino acid sequences of the *Ambiguiviridae* family genomes identified in this study (highlighted in red), representative members of the *Ambiguiviridae and Tombusviridae* families available in GenBank and ICTV-validated genomes. Node support was assessed using 1000 bootstrap replicates.

#### Alphambiguiviruses

Mycotombusviruses which are phylogenetically related to plant-infecting tombusviruses (Yang *et al*. 2024) were recently accommodated by the ICTV to a new family named *Ambiguiviridae* (https://ictv.global/taxonomy/taxondetails?taxnode_id=202521525&taxon_name=Ambiguiviridae). Ambiguiviruses typically possess a (+)ssRNA genome ranging from 2.6 to 5.5 kb, encoding two ORFs: one for a hypothetical protein of unknown function (P1) and another (P2) for a RdRp. A distinctive feature of this group is the presence of a ‘GDN’ catalytic motif in the RdRp, replacing the canonical ‘GDD’ motif found in most (+)ssRNA viruses, including tombusviruses. Three ambiguivirus viral genomes: Pyricularia oryzae RNA virus 1-1 (PoRV1-1, 3245 nt), PoRV1-2 (3248 nt) and PoRV1-3 (3245 nt) were recovered from *P. oryzae* isolates CH1184, CH1889 and CH2061, respectively (Table 1 and Supplementary Table 4). Each genome contained two in-frame ORFs separated by a 130-nucleotide intergenic region, and is flanked by 5ʹ-UTRs of 386–387 nt and 3ʹ-UTRs of 448–450 nt (Supplementary Figure 4A). ORF1 encoded a 265 aa hypothetical protein with unknown function (P1), sharing 96.6%-98.5% amino acid identity to the corresponding Magnaporthe oryzae RNA virus P1 protein (accession AJA41111, species *Alphambiguivirus magnaporthensis*), while ORF2 encoded a 494 aa RdRp featuring the metal-binding ‘GDN’ triplet in motif C (Supplementary Figure 4B). The genomic sequences of PoRV1-1, PoRV1-2 and PoRV1-3 shared 93.1-95% nucleotide identity with Magnaporthe oryzae RNA virus (accession KP174727) belonging to the species *Alphambiguivirus magnaporthensis*, a founding species of the *Alphambiguivirus* genus (*Ambiguiviridae* family). In addition, the RdRp proteins of PoRV1-1, PoRV1-2 and PoRV1-3 ORF2 shared 97.8-99% aa identity with the Magnaporthe oryzae RNA virus protein (accession AJA41112, *Alphambiguivirus magnaporthensis*) (Table 2), while the predicted P1-P2 readthrough protein sequences of these three viruses exhibited 97.3-98.6% aa identity with the P1-P2 readthrough protein sequences of Magnaporthe oryzae RNA virus (accession KP174727). Given the species demarcation threshold (<78% P1-P2 aa identity) for this family, PoRV1-1, PoRV1-2 and PoRV1-3 should tentatively belong to *Alphambiguivirus magnaporthensis* species. Phylogenetic analysis based on the RdRp and P1-P2 aa sequences (Figure 2C and Supplementary Figure 4C) confirmed that PoRV1-1, PoRV1-2 and PoRV1-3 cluster within the *Alphambiguvirus* genus.

### Negative-sense single-stranded RNA virus

#### Mymonaviruses

Two bipartite viral genomes, named Pyricularia oryzae mymonavirus 1-1 (PoMV1-1) and PoMV1-2, each composed by two ssRNA segments, were recovered from *P. oryzae* isolates CH1184 and CH2061, respectively (Table 1 and Supplementary Table 4). These two viral isolates shared 99.6% nucleotide identity across both genomic segments. The first segment (RNA1), measuring 4,442 nt (PoMV1-1) and 4,443 nt (PoMV1-2), encoded five putative non-overlapping ORFs, including a nucleoprotein (NP) and four hypothetical proteins of unknown function (Supplementary Figure 5a-A). Multiple sequence alignment of the putative gene-junction sequences between ORFs revealed a semi-conserved sequence motif, 3′- U/C)(UUA)(G/A)(G/C/A)AAAAA(C/A)C-5′(Supplementary Figure 5a-B), likely corresponding to conserved sequences characteristic of the order *Mononegavirales* (Jiāng *et al*. 2022). RNA1 shared the highest nucleotide identity (80.4%) with Magnaporthe oryzae mymonavirus 1 isolate NJ39 (accession OL415836).

The second segment (RNA2), approximately 6,081 nt (PoMV1-1) and 6,085 nt (PoMV1-2) in length, encoded RdRp proteins of 1,945 aa, featuring a catalytic domain L and a mRNA-capping region V domain containing the conserved motif GxxTx(n)HR, which is essential for mRNA cap formation (Supplementary Figure 5a-A). The RdRp protein sequences of PoMV1-1 and PoMV1-2 (RNA2) shared 95.4%–95.7% aa identity with the corresponding L protein of Magnaporthe oryzae mymonavirus 1 isolate NJ39 (accession UYO08140) and 90.9%-90.7% aa identity with Magnaporthe oryzae mononegaambi virus 1 (accession QVU39967) belonging to the species *Penicillimonavirus magnaporthe*.

Among *P. oryzae* mymonavirus isolates with both RNA1 and RNA2 segments deposited in GenBank, only strains NJ19 and RLB18 possess sequences for both segments. In contrast, isolate NJ39 carries a single 10,515 nt RNA segment encoding both genes (accession OL415836). For isolate RLB18 (accession ON791637 [RNA1] and accession ON791636 [RdRp_RNA2]), the two partial segments of 3,681 nt and 4,224 nt were obtained by Illumina sequencing and remain incomplete (He *et al*. 2022). In this study, we identified two distinct RNA segments, with no sequencing reads spanning their junction. PCR amplification using a primer pair designed across the putative junction (Supplementary Table 2) failed to yield a product consistent with a monopartite genome, supporting a bipartite organization. Furthermore, RACE-PCR successfully recovered the terminal sequences of both RNA segments (Supplementary Figure 5b).

Phylogenetic analysis generated from the amino acid sequences of the RdRp of mymonaviruses confirmed that PoMV1-1 and PoMV1-2 were closely related to Magnaporthe oryzae mononegaambi virus 1 (accession QVU39967.1, species *Penicillimonavirus magnaporthe*) (Figure 3A). While PoMV1-1 and PoMV1-2 differ by <30% in nucleoprotein amino acid sequence and <30% in complete genome nucleotide sequence from *Penicillimonavirus magnaporthe*, we tentatively propose that these two bipartite isolates (PoMV1-1 and PoMV1-2) belong to this species.

**Figure 3.** **(A)** A maximum-likelihood phylogenetic tree was constructed based on the RdRp amino acid sequences of the *Mymonaviridae* family genomes identified in this study (highlighted in red) and representative members of the *Mymonaviridae* family available in GenBank. The analysis includes ICTV-validated genomes of different genera within the family. Node support was assessed using 1000 bootstrap replicates. **(B)** A maximum-likelihood phylogenetic tree was constructed based on the RdRp amino acid sequences of the *Partitiviridae* family genomes identified in this study (highlighted in red) and representative members of the *Partitiviridae* family available in GenBank. The analysis includes ICTV-validated genomes of different genera within the family. Node support was assessed using 1000 bootstrap replicates. **(C)** A maximum-likelihood phylogenetic tree was constructed based on the RdRp amino acid sequences of the *Polymycoviridae* family genomes identified in this study (highlighted in red), representative members of the *Polymycoviridae* family available in GenBank and ICTV-validated genomes of the *Polymycoviridae* family. The analysis includes ICTV-validated genomes of different genera within the family. Node support was assessed using 1000 bootstrap replicates.

### Double-stranded RNA virus

#### Partitiviruses

A viral genome, Pyricularia oryzae partitivirus 2 (PoPV2), was recovered from *P. oryzae* isolate CH2054. Its putative capsid protein gene, carried by RNA2 (1,491 nt, 420 aa), shared 99.6% nucleotide identity with Magnaporthe oryzae partitivirus 1 strain NJ20 segment 2 (accession KX119173) and 99.3 % nucleotide identity with Magnaporthe oryzae partitivirus 2 (accession KX981864) (Table 1 and Supplementary Table 4). The *RdRp* gene contained by RNA1 (1,763 nt, 539 aa) shared 99.7% nucleotide identity with both Magnaporthe oryzae partitivirus 2 (accession KX981863) and Magnaporthe oryzae partitivirus 2A strain RLB18 (accession ON791632) as well as 98.1% with Magnaporthe oryzae partitivirus 1 strain NJ20 segment 1 (accession KX119172) (Table 1), these three partitiviruses putatively belonging to the genus *Gammapartitivirus*. The RdRp and CP sequences of PoPV2 shared only 69.5% and 63% amino acid identity, respectively, with the closest ICTV-recognized reference virus Penicillium stoloniferum virus F (accession AAU95758) belonging to the species *Gammapartitivirus effepenicillii*, which is below the established species demarcation threshold (<90% and <80% aa sequence identity for RdRp and CP, respectively) (Table2). Consistently, phylogenetic analysis based on RdRp aa sequences (Figure 3B) confirmed that PoPV2 clustered with Magnaporthe oryzae partitivirus 2A (accession UVX28891), Penicillium stoloniferum virus F (accession AAU95758) and other putative members of the *Gammapartitivirus* genus, supporting the classification of PoPV2 as a member of a novel species within this genus.

#### Polymycoviruses

A multi-segmented viral genome, Pyricularia oryzae polymycovirus 1 (PoPmV1), composed of five dsRNA segments ranging from 1,317 to 2,397 bp (Supplementary Figure 6A) was recovered from *P. oryzae* isolate CH1184 (Table 1 and Supplementary Table 4). DsRNA1 encoded a RdRp (766 aa), while the remaining segments (dsRNA2–dsRNA5) encoded hypothetical proteins (Supplementary Figure 6A). Segments dsRNA1–dsRNA4 of PoPmV1 shared 86.7–92.3% nucleotide identity with the four corresponding segments of Magnaporthe oryzae polymycovirus 1_isolate TM02 (dsRNA1: accession MH231406, dsRNA2: accession MH231407, dsRNA3: accession MH231408 and dsRNA4: accession MH231409) belonging to the species *Multimycovirus magnaporyzae*. The RdRp protein of PoPmV1 shared 97.4% aa identity with the same reference virus Magnaporthe oryzae polymycovirus 1 (accession QAU09249) (Table 2).

DsRNA2 encoded a putative serine protease (697 aa) sharing 98.1% identity with Magnaporthe oryzae polymycovirus 1 (accession QAU09250), while dsRNA3 encoded a putative methyltransferase (612 aa) exhibiting 95.9% identity with Magnaporthe oryzae polymycovirus 1 (accession QAU09251). DsRNA4 contained two ORFs: ORF4a encoding a PASr protein (263 aa) sharing 96.2% aa identity with Magnaporthe oryzae polymycovirus 1 (accession QAU09252) and ORF4b, encoding a 33 aa protein of unknown function. The fifth segment, dsRNA5, validated by RT-PCR, encoded a 392 aa protein of unknown function, sharing 46% and 44.4% (coverage 19% and 22%) aa identity with ORF7 of Beauveria bassiana polymycovirus 2 (accession CUS18601) and ORF6 of Beauveria bassiana polymycovirus 3 (accession CAD7829828), respectively. The 5′ termini of all dsRNA segments shared a conserved motif, (GAA(C/T)TTAAGAGTTTTTCT(A/G/C)C(GA)C(A/T)(A/T)G(C/T), while the 3′ termini uniformly ended with “ATTT” (Supplementary Figure 6B-6C). The conservation of terminal motifs confirmed that RNA5 belongs to the same viral genome.

A phylogenetic analysis based on RdRp aa sequences (Figure 3C), including PoPmV1 and representative members of the *Polymycoviridae* family, showed that PoPmV1 clustered with members of the *Multimycovirus* genus. According to ICTV species demarcation criteria (≤ 70% aa RdRp identity and clustering within a highly supported monophyletic clade with members of the same genus), our results suggest that PoPmV1 belongs to *Multimycovirus magnaporyzae* species. Notably, however, the MoPmV1 isolate contains five genomic segments whereas the exemplar isolate of Magnaporthe oryzae polymycovirus 1_ isolate TM02 (Zheng *et al*. 2025) carries only four.

## Discussion

Here, we present the MADAM approach, which integrates multiple technical steps that have been independently employed in other protocols. By combining these modular components, we have developed a highly efficient mycovirus detection method that yields an exceptionally high proportion of viral reads. The MADAM approach leverages dsRNA molecules, which serves as a highly reliable molecular marker for virus detection across animal, plant, and fungal hosts (Roossinck *et al*. 2010; Marais *et al*. 2018b; Crabtree *et al*. 2019; Decker *et al*. 2019; Gaafar & Ziebell 2020; Javaran *et al*. 2023). Specifically, by selectively targeting dsRNA, this method minimizes host nucleic acid amplification, thereby enhancing sequencing depth and enabling the recovery of more complete viral genomes. Consequently, this selective enrichment is critical for both sensitive viral detection and the discovery of novel viruses. The MADAM approach exploits anti-dsRNA mAbs, which eliminate the requirement for target nucleotide sequence recognition, thereby enabling universal viral detection. This specificity arises from a structure-dependent interaction: mAb engages its dsRNA target through a concerted action of its heavy and light chain complementarity-determining regions. These regions collaboratively trace the minor groove of the dsRNA helix, thereby conferring the antibody’s high-affinity and sequence-independent binding properties (Bou-Nader *et al*. 2025). Monoclonal antibody–mediated dsRNA enrichment has proven effective in animal and plant systems (O’Brien *et al*. 2015; Blouin *et al*. 2016; Decker *et al*. 2019) and our results further demonstrate its applicability to fungal samples.

The MADAM approach incorporates random deep sequencing, a method demonstrated to be highly effective for the comprehensive detection and characterization of both known and novel RNA viruses (Gaafar & Ziebell 2020; Young *et al*. 2021; Javaran *et al*. 2023; Pichler *et al*. 2023). This process adheres closely to the Oxford Nanopore Direct cDNA Sequencing Kit protocol (PCS109), which employs random reverse primers—rather than poly(T) primers (VNP), in a strand-switching reaction, followed by second-strand synthesis and PCR amplification. In our study, the inclusion of a second-strand synthesis step prior to PCR amplification was essential to generate sufficient material for library preparation.

Notably, some previous studies have omitted either the second-strand synthesis step, PCR amplification, or both (Young *et al*. 2021; Javaran *et al*. 2023; Pichler *et al*. 2023; Fall *et al*. 2025). As a result, direct comparisons of viral read outputs across studies remain challenging. Variations in viral read abundance are likely influenced primarily by sample type and initial RNA concentration, both of which can significantly affect dsRNA yield. Additionally, methodological differences, particularly in dsRNA enrichment protocols and RNA virus amplification strategies, are expected to contribute to observed discrepancies in viral read recovery between studies.

Finally, the MADAM approach employed Oxford Nanopore Flongle sequencing, a platform validated for rapid genome detection in clinical, plant, and veterinary diagnostics (Young *et al*. 2021; Javaran *et al*. 2023; Pichler *et al*. 2023; Fall *et al*. 2025). Its advantages, rapid turnaround, cost-efficiency, and single-sample compatibility, offset higher per-base error rates, which we mitigated via reference-based assembly and high-coverage consensus sequences (Ben Chehida *et al*. 2021). Long reads enabled de novo assembly and novel virus discovery, while barcoded multiplexing reduced per-sample costs. For terminal genome characterization, poly(A)-targeted primers associated with PCR amplification and ONT sequencing were used, avoiding laborious cloning of PCR products to account for tail-length variability.

The MADAM approach was initially benchmarked against the VANA approach, the latter being recognized for its ability to detect both RNA and DNA viruses (Moubset *et al*. 2022). While both metagenomic methods identified the same set of mycoviruses, the viral read recovery rate (0.6–14.1%) and the sequencing coverage and depth provided by the VANA approach proved insufficient for reconstructing all known and novel viral genomes. In contrast, the MADAM approach achieved superior qualitative and quantitative performance, enabling comprehensive genome reconstruction. Specifically, the MADAM approach achieved a high viral read recovery rate (46.9–72.7%), consistent with plant virus study reporting 31.4-74.4% recovery (Blouin *et al*. 2016). However, read distribution among co-infecting viruses varied markedly, likely reflecting differences in replication dynamics and intra-host viral titers. The method produced near-complete to complete viral genomes, with average sequencing depths ranging from 18X to 78,195X across all viral genomes. Fifteen complete genomes shared between 80.4–99.7% nucleotide identity to previously characterized *P. oryzae* viruses. Notably, the MADAM approach enabled the high-coverage detection of previously undescribed viral genomes, including PoBV16 (3,557X average depth), a new member of the *Botourmiaviridae* and PoPmV1- dsRNA5 (3,417X average depth), a fifth genomic segment of the polymycovirus PoPmV1, as well as the first representative of the *Deltaormycoviridae* family in *P. oryzae*. For novel sequences exhibiting <65% nucleotide similarity and <37% coverage to top BLASTn hits, assembly relied on average read lengths of ∼400 nt, with a subset of longer reads (800–1,800 nt, comprising ∼2% of total reads per virus), which ensured robust *de novo* reconstruction. All newly identified genomes and terminal sequences were independently validated through RT-PCR using targeted primers and overlapping amplicons.

The MADAM approach also enabled the high-coverage detection of previously described viral genomes, including PoBV14, which shares 97.6% aa identity with Magnaporthe oryzae botourmiavirus 14 (accession XNZ83248) that was proposed to belong to the genus *Gammascleroulivirus* (Li *et al*. 2025). However, this taxonomic assignment has not yet been officially recognized by ICTV. Our results suggest that this proposed classification may not align with current ICTV demarcation criteria. Specifically, the RdRp amino acid sequences of PoBV14 and Magnaporthe oryzae botourmiavirus 14 share less than 53% identity with the closest recognized members of the *Botourmiaviridae* family (Table 2). This value falls substantially below the 70% amino acid identity threshold proposed for genus-level demarcation. Thus, PoBV14 and Maganoporthe oryzae botourmiavirus 14 may instead represent members of a previously undescribed genus within the family *Botourmiaviridae*.

Furthermore, a fifth genomic segment, dsRNA5, was identified in the polymycovirus PoPmV1, although its function remains unknown. Investigating additional *Pyricularia oryzae* isolates will be important to determine whether this segment is conserved and to assess its potential. functional role. In particular, characterization of the encoded viral ORF(s), following the approach used to study ORF5 in *Beauveria bassiana* (Shi *et al*. 2025), could help elucidate its biological role and determine whether it contributes to stress tolerance and hypervirulence, as reported for P5 through its interactions with host proteins. In addition, the proline-alanine-serine-rich protein (PASrp) encoded by MoPmV1 dsRNA4a may function as a putative structural component involved in the unconventional coating of the dsRNA genome, as reported for other polymycoviruses (Zheng *et al*. 2025). Notably, this segment shares 43.6% identity with ORF5 of Pseudopestalotiopsis camelliae-sinensis polymycovirus 1 (accession XHV10074.1), which has been proposed to mediate filamentous virion encapsidation based on experimental observations in Phyllosticta capitalensis polymycoviruses 1 and 2 and Botryosphaeria dothidea RNA virus 1 (BdRV1) (Han *et al*. 2024). A similar role in filamentous virion encapsidation may also be hypothesized for the MoPmV1 and PoPmV1 PASrp. The deltaormycoviruses PoDOmV1-1 and PoDOmV1-2 were markedly underrepresented in both VANA and MADAM analyses, accounting for less than 1% of total viral reads. This highlights the need for specific enrichment to achieve adequate sequencing depth for reliable detection. Although the overall genome coverage for both segments was moderate to high (103X–733X average depth) with MADAM method, their terminal regions remained incomplete and were resolved experimentally using segment-specific primers. BLASTn analysis revealed 69.4–69.6% and 75.4–75.7% nucleotide identity for each segment, respectively, with the closest BLASTn match being Verticillium dahliae ormycovirus 1. No closely related equivalents have been previously reported in *P. oryzae*.

Similarly, RNA2 of the mymonaviruses PoMV1-1 and PoMV1-2 (∼6,000 nt) exhibited reduced sequencing coverage, requiring RT-PCR validation and completion with segment-specific primers. *Mymonaviridae* members possess (-)ssRNA genomes, and dsRNA-targeting monoclonal antibodies show low capture efficiency for such viruses (O’Brien *et al*. 2015). While negative-sense RNA viruses were initially believed to lack dsRNA intermediates, recent viromic studies have demonstrated that (-)ssRNA viruses do produce low amounts of dsRNA during replication detectable via mAb (Son *et al*. 2015). Consistently, two other studies have reported detection of a putative emaravirus (another (-)ssRNA virus) using dsRNA enrichment strategies (Blouin *et al*. 2016; Rabbidge *et al*. 2021). Our findings further confirm that (-)ssRNA viruses can be detected with 2G4 mAb-based approaches, albeit with lower read abundance. It is noteworthy that PoMV1-2 RNA segments 1 and 2, detected using the VANA method, yielded a higher number of sequencing reads despite the overall lower total viral read count compared to the MADAM approach. This discrepancy may be attributed to the fact that genomic RNAs of many negative-sense single-stranded RNA viruses are tightly associated with nucleoproteins during replication, potentially limiting their accessibility for dsRNA-based enrichment (Ruigrok *et al*. 2026). However, sequencing depth patterns suggested a bipartite genome organization for this *Mymonaviridae* member, which was subsequently validated by the absence of RT-PCR amplification across segment junction. The terminal regions of each segment were validated experimentally using segment-specific primers (Supplementary Figure 5b). To date, bipartite genomes have not been reported in the *Mymonaviridae* family, except for Plasmopara viticola lesion-associated mononegaambi virus 3, first described by Morán et al. (Morán *et al*. 2023) and later confirmed by Pagnoni et al. (Pagnoni *et al*. 2023) in Trichoderma harzianum mononegavirus 1 (ThMV1).

While no DNA viruses have yet been reported to date in *P. oryzae*, a similar study by Rott et al. (Rott *et al*. 2024), which also employed monoclonal antibodies (mAb J2), successfully detected DNA viruses, as further confirmed by immunofluorescence analysis (Son *et al*. 2015). However, the use of dsRNA sequencing combined with anti-dsRNA monoclonal antibodies for the detection of DNA viruses still requires further validation.

## Conclusion

The MADAM approach integrates multiple technical modules previously employed independently in other protocols. Specifically, it combines dsRNA enrichment via monoclonal antibodies (mAbs) with adapter-modified random octamer priming in a strand-switching reaction, followed by Oxford Nanopore MinION sequencing, to enable high-resolution detection and characterization of RNA mycoviruses. By incorporating full-length genome sequencing, this workflow substantially enhances viral sequence diversity assessment, thereby advancing our understanding of the global mycovirosphere. Moreover, its robustness and scalability make it highly valuable for routine diagnostic and surveillance applications.

## Acknowledgements

We are grateful to Jody Hobson-Peters for her generous assistance in providing the dsRNA antibodies used in this study.

## Data accessibility statement

The nucleotide sequences used in the phylogenetic analyses are publicly available in the GenBank database (accession numbers PZ324806 to PZ324813 and PZ356821 to PZ356842). The cleaned sequence read data from the Nanopore runs are available from the NCBI Sequence Read Archive, under the Bioproject accession number PRJNA1465638 (Biosample accession numbers SAMN59750546, SAMN59750547, SAMN59750548, SAMN59750549). Scripts were deposited online at: https://zenodo.org/records/20205917).

