## Supplementary Material for "Monoclonal anti-dsRNA antibody-based metagenomics (MADAM) reveal *Pyricularia oryzae* mycovirome"

| *P. oryzae* strain | Year | Country | Province | Location | Landrace vernacular name |
|---|---|---|---|---|---|
| CH1184 | 2009 | China | Yunnan | Xinjie | Ai Zhe Gu |
| CH1889 | 2015 | China | Yunnan | Jingkou | Xiaogu |
| CH2054 | 2016 | China | Yunnan | Gingko | HY3 |
| CH2061 | 2016 | China | Yunnan | Da Yu Tang | HY3 |

Supplementary Table 1 : List of *P. oryzae* strains, including the vernacular variety names and the villages where samples were collected

| Objectives | Primer name | 5'–3' nt sequence | Purpose |
|---|---|---|---|
| Primers used for cDNA-PCR and for characterizing all 5'-3' extremities | VNP-8N | 5'-/5Phosp/ACTTGCCTGTCGCTCTATCTTCN8-3' | RT primer and second strand synthesis |
| | SSP | TTTCTGTTGGTGCTGATATTGCTGCCATTACGGCCmGmGmG | RT primer |
| | PR2 | TTTCTGTTGGTGCTGATATTGC | Second strand synthesis and PCR |
| | VNP | ACTTGCCTGTCGCTCTATCTTCTTTTTTTTTTTTTTTTV | RT primer after poly A tailing |
| | 3580R | ACTTGCCTGTCGCTCTATCTTC | PCR |
| PCR jonction between two segment RNA1 and RNA2 of mymonavirus | CH2061-Mymona1-3874F | GGAATGACGCCAAGAGCACC | 4 PCRs/combination between Forward and Reverse/ Size PCRs products expected between 373 and 1106 bp |
| | CH2061-Mymona1-4295F | CTCCGTAGTACTCAGGCCTC | |
| | CH2061-Mymona2-562R | ATGCTGTTCTATACGCGGAC | |
| | CH2061-Mymona2-222R | CGTTGAGATGAGAGTCATCG | |
| Mymonavirus 5' and 3' extremities | CH2061_Mymona1_498R | CTCAATTCACCAGTCTCACG | PCR with 3580R primer - 3' extremity PoMV1-1/PoMV1-2 RNA1 |
| | CH2061_Mymona1_3798F | CATGCTCGCTACTCGAATGT | PCR with 3580R primer - 5' extremity PoMV1-1/PoMV1-2 RNA1 |
| | CH2061_Mymona2_562R | ATGCTGTTCTATACGCGGAC | PCR with 3580R primer - 3' extremity PoMV1-1/PoMV1-2 RNA2 |
| | CH2061-Mymona2_5521F | ACCAATGTCCACCACTGCAC | PCR with 3580R primer - 5' extremity PoMV1-2 RNA2 |
| | CH2061_Mymona2 5424F | TCACTTGAGCTTGCATCACG | PCR with 3580R primer - 5' extremity PoMV1-1 RNA2 |
| | CH1184_Mymona2_5475F | CTCATTCGCACTCAGCTATAGAGGT | PCR with 3580R primer - 5' extremity PoMV1-1 RNA2 |
| CH2061 - Mymonavirus segment2 internal gap | CH2061_Mymona2_3467F | CATCACGTTCGCTCATGAAG | Sanger sequencing PCR product ~ 555bp |
| | CH2061_Mymona2_4022R | GCATAATCACTTGTGGCGAC | |
| Deltaormycovirus 5' and 3' extremities | CH1889_Ormyco1_3'-2593F | ATACCGAACGTACTAGCAGC | PCR with 3580R primer - 3' extremity PoDOmV1-1 RNA1 |
| | CH1889_Ormyco1_5'_653R | AGTATGTGGCCATACGACTG | PCR with 3580R primer - 5' extremity PoDOmV1-1 RNA1 |
| | CH1889-Ormyco1_2620F | AAGCTTCCAGTAGACTTTAGAGTG | PCR with 3580R primer - 3' extremity PoDOmV1-1 RNA1 |
| | CH1889_Ormyco2-1015F | GTAGACCAGGAAGTGATCTC | PCR with 3580R primer - 3' extremity PoDOmV1-1 RNA2 |
| | CH1889_Ormyco2_656R | CGGAGCGATGATTGTATGTC | PCR with 3580R primer - 5' extremity PoDOmV1-1 RNA2 |
| | CH1889_Ormyco1_3'_2682F | CAGTAGAACTGGTAGACTTCCTAT | PCR with 3580R primer - 3' extremity PoDOmV1-1 RNA1 |
| CH2061- Botourmiavirus 16 (PoBV16) - 5' and 3' extremities and internal PCR | CH2061_ourmia_like_virus_653F | GACCAGTTGCTCCTCCACGA | internal PCR product_PoBV16 |
| | CH2061_ourmia_like_virus__1326R | GGAAGACGCTCGTGGATCCT | |
| | CH2061_ourmia_like_virus__1221F | AACAACGCTCGTGGCCTAAC | internal PCR product_PoBV16 |
| | CH2061_ourmia_like_virus__1943R | CCTACTCCGCCGTTGTGCTT | |
| | CH2061_ourmia_like_virus__47F | TGTAGTAGGCCTGTACGCTG | internal PCR product_PoBV16 |
| | CH2061_ourmia_like_virus__757R | TTCTACATCCGCAAGAGCCT | |
| | CH2061_ourmia_like_virus__1863F | CCATCAACAACGGTAGAGAC | internal PCR product_PoBV16 |
| | CH2061_ourmia_like_virus__2493R | TCCTGATCCTGAACCTTGAC | |
| | CH2061_ourmia_like_virus_-672R | TCGTGGAGGAGCAACTGGTC | PCR with 3580R primer -5' extremity PoBV16 |
| | CH2061_ourmia_like_virus__275R | GCGTACGAACCGTGAACACC | PCR with 3580R primer -5' extremity PoBV16 |
| | CH2061_ourmia_like_virus__2131F | CCATACGGCGAACGTAGTCC | PCR with 3580R primer -3' extremity PoBV16 |
| CH1184 - Polymycovirus RNA4 - 5' and 3' extremities | CH1184_Polymycovirus_RNA4_810F | GCGTCTACTTCATTCGTGAG | PCR with 3580R primer -5' extremity PoPmV1 RNA4 |
| | CH1184_Polymycovirus_RNA4_862F | CGAGAAGGTGAAGTTCGACG | PCR with 3580R primer -5' extremity PoPmV1 RNA4 |
| | CH1184_Polymycovirus_RNA4_522R | CGTGAAGAGCCGGAGATACG | PCR with 3580R primer -3' extremity PoPmV1 RNA4 |
| | CH1184_Polymycovirus_RNA4_703R | ACAGCGGTAACGCAATGGAC | PCR with 3580R primer -3' extremity PoPmV1 RNA4 |
| CH1184 - Polymycovirus RNA5 - 5' and 3' extremities and internal PCR | CH1184_Polymycovirus_RNA5_1146R | GAGTTGCTGAGCGAGACGAT | Internal PCR product |
| | CH1184_Polymycovirus_RNA5_394F | GGCACACAACTAGCACCGTC | |
| | CH1184_Polymycovirus_RNA5_868F | CTCGAGTGCACGAATCATCC | PCR with 3580R primer -5' extremity PoPmV1 RNA5 |
| | CH1184_Polymycovirus_RNA5_832F | GATCATGGTTGGAGCACGGT | PCR with 3580R primer -5' extremity PoPmV1 RNA5 |
| | CH1184_Polymycovirus_RNA5_882R | ATTCGTGCACTCGAGGAGAC | PCR with 3580R primer -3' extremity PoPmV1 RNA5 |
| | CH1184_Polymycovirus_RNA5_1004R | GTCAGTGAGACTAATCGTCG | PCR with 3580R primer -3' extremity PoPmV1 RNA5 |

Supplementary Table 2: RT-PCR, PCR, and sequencing primers (5'–3')

| Strains | Total reads number after QC | Viral reads (% of total reads) | Virus name abbreviation(s) | Virus names | Segment | Predicted Protein Encoded | Mapping reads (% viral reads) | Average deep sequencing coverage | Coverage (%) (a) |
|---|---|---|---|---|---|---|---|---|---|
| CH2054 | 110624 | 649 (0.58%) | PoPV2 | Pyricularia oryzae partitivirus | dsRNA1 | RdRp | 132 (20.3%) | 9.4 | 90.4 |
| | | | | | dsRNA2 | CP | 392 (60%) | 32.3 | 93.4 |
| CH2061 | 245 027 | 34622 (14.1%) | PoBV14 | Pyricularia oryzae botourmiavirus 14 | | RdRp | 202 (0.58%) | 11.1 | 66.0 |
| | | | PoBV4 | Pyricularia oryzae botourmiavirus 4 | | RdRp | 477 (1.38%) | 23.6 | 76.4 |
| | | | PoBV6-2 | Pyricularia oryzae botourmiavirus 6-2 | | RdRp | 122 (0.35%) | 10.9 | 61.7 |
| | | | PoBV7 | Pyricularia oryzae botourmiavirus 7 | | RdRp | 256 (0.74%) | 17 | 75.5 |
| | | | PoBV16 | Pyricularia oryzae botourmiavirus 16 | | RdRp | 217 (0.63%) | 11.1 | 59.3 |
| | | | PoMV1-2 | Pyricularia oryzae mymonavirus 1-2 | RNA1 | HP1-HP5 | 28983 (83.7%) | 863 | 99.6 |
| | | | | | RNA2 | RdRp | 2017 (5.8%) | 42.5 | 99.1 |
| | | | PoDOmV1-2 | Pyricularia oryzae deltaormycovirus 1-2 | RNA1 | RdRp | 16 (0.05%) | 0.6 | 30.3 |
| | | | | | RNA2 | HP | 46 (0.13%) | 3.7 | 58.2 |
| | | | PoRV1-3 | Pyricularia oryzae RNA virus 1-3 | | HP/RdRp | 850 (2.5%) | 38.2 | 96.3 |

(a) Genome coverage relative to the total viral genome size obtained by Nanopore sequencing (Table 1-Supplementary Table 3)

Supplementary Table 3: Information on the assembled reads of mycoviruses obtained from VANA sequencing method

| Strains | Putative family/ genus virus | | GenBank ID | Virus names | Virus name abbreviation(s) | Segment | Predicted Protein Encoded | Sequence size | Closest relative virus (BLASTn) | % id nt sequence | Query coverage | E-value | Closest relative virus (BLASTp) | % id CDS aa sequence | Query coverage | E-value |
|---|---|---|---|---|---|---|---|---|---|---|---|---|---|---|---|---|
| CH1184 | *Ambiguiviridae/Alphambiguivirus* | | PZ356821 | Pyricularia oryzae RNA virus 1-1 | PoRV1-1 | | HP/RdRp | 3245 | KP174727 | 93.1 | 100% | 0.0 | AJA41111/AJA41112 | 96.6/97.7 | 100.0 | 0.0 |
| | *Polymycoviridae/Multimycovirus* | | PZ356834 | Pyricularia oryzae polymycovirus 1 | PoPmV1 | dsRNA1 | RdRp | 2397 | MH231406 | 92.3 | 100% | 0.0 | QAU09249 | 97.4 | 100.0 | 0.0 |
| | | | PZ356835 | | | dsRNA2 | HP | 2229 | MH231407 | 90.7 | 100% | 0.0 | QAU09250 | 98.1 | 100.0 | 0.0 |
| | | | PZ356836 | | | dsRNA3 | HP | 1959 | MH231408 | 86.7 | 100% | 0.0 | QAU09251 | 95.9 | 100.0 | 0.0 |
| | | | PZ356837 | | | dsRNA4 | HP | 1317[a] | MH231409 | 89.6 | 100% | 0.0 | QAU09252 | 96.2 | 100.0 | 0.0 |
| | | | PZ356838 | | | dsRNA5 | HP | 1388[a][b] | - | - | - | - | CUS18601 | 46.0 | 19.0 | 3.0E-06 |
| | *Mymonaviridae/Penicillimonavirus* | | PZ356839 | Pyricularia oryzae mymonavirus 1-1 | PoMV1-1 | RNA1 | HP1-HP5 | 4442[a] | OL415836 | 80.4 | 100% | 0.0 | UYO08135/QVU39968/ UYO08137/ UYO08138/UYO08139 | 90.3/98.5/91.7/ 88.6/84.7 | 100.0/89.0 (HP4) | $8e^{-149}$/0.0/$6e^{-128}$/ $1e^{-135}$/$2e^{-105}$ |
| | | | PZ356840 | | | RNA2 | RdRp | 6081[a] | OL415836 | 86.5 | 100% | 0.0 | QVU39974 | 95.9 | 100.0 | 0.0 |
| CH1889 | *Botourmiaviridae* | */Magoulouvirus* | PZ324806 | Pyricularia oryzae botourmiavirus 2 | PoBV2 | | RdRp | 2255 | MW117114 | 94.6 | 100% | 0.0 | UVX28883 | 96.9 | 100.0 | 0.0 |
| | | */Scleroulivirus* | PZ324807 | Pyricularia oryzae botourmiavirus 3 | PoBV3 | | RdRp | 2818 | LC413503 | 94.4 | 91% | 0.0 | BBF90578 | 96.8 | 100.0 | 0.0 |
| | | */Gammascleroulivirus* | PZ324808 | Pyricularia oryzae botourmiavirus 6-1 | PoBV6-1 | | RdRp | 2370 | MW752191 | 92 | 99% | 0.0 | QVU39993 | 95.7 | 100.0 | 0.0 |
| | *Splipalmiviridae/Delepalmivirus* | | PZ356830 | Pyricularia oryzae splipalmivirus 1 | PoSpV1 | RNA1 | RdRp | 2455 | LC553711 | 96.4 | 99% | 0.0 | BCH36656 | 99.0 | 100.0 | 0.0 |
| | | | PZ356831 | | | RNA2 | RdRp | 2480 | LC553710 | 96.8 | 99% | 0.0 | BCH36655 | 98.9 | 100.0 | 0.0 |
| | | | PZ356832 | | | RNA3 | HP | 1980 | LC553712 | 97.7 | 99% | 0.0 | BCH36657 | 98.4 | 100.0 | 0.0 |
| | | | PZ356833 | | | RNA4 | HP | 1226 | LC553713 | 98.2 | 99% | 0.0 | BCH36658 | 99.1 | 100.0 | 0.0 |
| | *Ambiguiviridae/Alphambiguivirus* | | PZ356822 | Pyricularia oryzae RNA virus 1-2 | PoRV1-2 | | HP/RdRp | 3248 | KP174727 | 93.5 | 100% | 0.0 | AJA41111/AJA41112 | 98.1/99.0 | 100.0 | 0.0 |
| | *Deltaormycoviridae/Bormycovirus* | | PZ356826 | Pyricularia oryzae deltaormycovirus 1-1 | PoDOmV1-1 | RNA1 | RdRp | 3357[a] | OR734290 | 69.6 | 96% | 0.0 | WPV08068 | 65.7 | 100.0 | 0.0 |
| | | | PZ356827 | | | RNA2 | HP | 1537[a] | OR734291 | 75.7 | 88% | 0.0 | WPV08069 | 74.4 | 96.0 | 0.0 |
| CH2054 | *Partitiviridae/Gammapartitivirus* | | PZ356824 | Pyricularia oryzae partitivirus 2 | PoPV2 | dsRNA1 | RdRp | 1763 | KX981863 | 99.7 | 100% | 0.0 | UVX28891 | 100.0 | 100.0 | 0.0 |
| | | | PZ356825 | | | dsRNA2 | CP | 1491 | KX119173 | 99.6 | 100% | 0.0 | APP18152 | 99.8 | 100.0 | 0.0 |
| CH2061 | *Botourmiaviridae* | /Unclassified | PZ324812 | Pyricularia oryzae botourmiavirus 14 | PoBV14 | | RdRp | 2322 | PQ732967 | 95.3 | 100% | 0.0 | UUW21043 | 98.0 | 100.0 | 0.0 |
| | | */Penoulivirus* | PZ324811 | Pyricularia oryzae botourmiavirus 4 | PoBV4 | | RdRp | 2544 | MK507958 | 97.0 | 98% | 0.0 | QDW80874 | 96.6 | 100.0 | 0.0 |
| | | */Gammascleroulivirus* | PZ324810 | Pyricularia oryzae botourmiavirus 6-2 | PoBV6-2 | | RdRp | 2366 | MN971591 | 91.3 | 100% | 0.0 | UUW21042 | 94.0 | 100.0 | 0.0 |
| | | */Epsilonscleroulivirus* | PZ324809 | Pyricularia oryzae botourmiavirus 7 | PoBV7 | | RdRp | 2321 | MN971592 | 94.8 | 100% | 0.0 | QLJ94431 | 96.2 | 100.0 | 0.0 |
| | | /Unclassified | PZ324813 | Pyricularia oryzae botourmiavirus 16 | PoBV16 | | RdRp | 2502[a][b] | MN565694 | 65.6 | 37% | 9E-24 | QIP68026 | 53.7 | 98.00 | 0.00 |
| | *Mymonaviridae/Penicillimonavirus* | | PZ356841 | Pyricularia oryzae mymonavirus 1-2 | PoMV1-2 | RNA1 | HP1-HP5 | 4443[a] | OL415836 | 80.4 | 100% | 0.0 | UYO08135/QVU39968/ UYO08137/ UYO08138/UYO08139 | 90.3/98.5/91.7/ 88.6/84.7 | 100.0/89.0 (HP4) | $8e^{-149}$/0.0/$6e^{-128}$/ $1e^{-135}$/$4e^{-105}$ |
| | | | PZ356842 | | | RNA2 | RdRp | 6085[a][b] | OL415836 | 86.3 | 100% | 0.0 | QVU39974 | 95.9 | 100.0 | 0.0 |
| | *Deltaormycoviridae/Bormycovirus* | | PZ356828 | Pyricularia oryzae deltaormycovirus 1-2 | PoDOmV1-2 | RNA1 | RdRp | 3357[a] | OR734290 | 69.4 | 96% | 0.0 | WPV08068 | 65.3 | 100.0 | 0.0 |
| | | | PZ356829 | | | RNA2 | HP | 1537[a] | OR734291 | 75.4 | 89% | 0.0 | WPV08069 | 74.6 | 96.0 | 0.0 |
| | *Ambiguiviridae/Alphambiguivirus* | | PZ356823 | Pyricularia oryzae RNA virus 1-3 | PoRV1-3 | | HP/RdRp | 3245 | KP174727 | 95.0 | 100% | 0.0 | AJA41111/AJA41112 | 98.5/98.6 | 100.0 | 0.0 |

(a) : Full-length sequence including both 5' and 3' ends obtained/validated using RACE sequencing.
(b) Gap filling was performed by PCR amplification with specific primers followed by Nanopore or Sanger sequencing.

Supplementary Table 4: Analysis of Nanopore-derived mycovirus contigs with BLASTn and BLASTp-based taxonomic assignment of assembled sequences

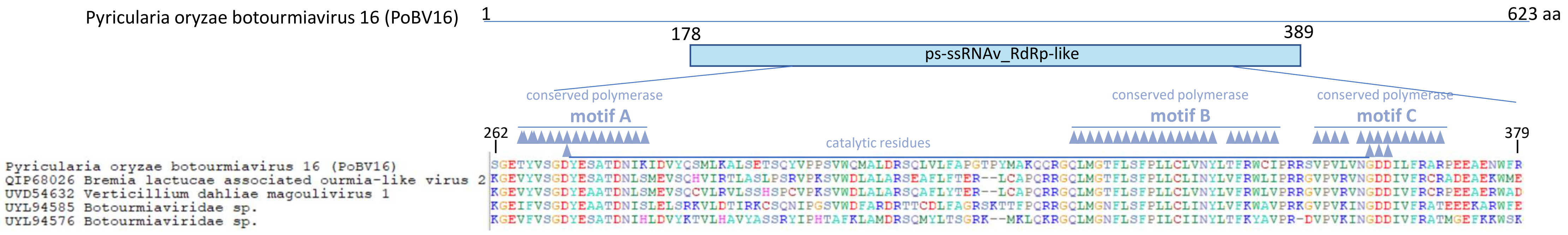


Supplementary Figure 1: Conserved motifs A, B, and C in PoBV16 RdRp, as annotated in NCBI Conserved Domain Database

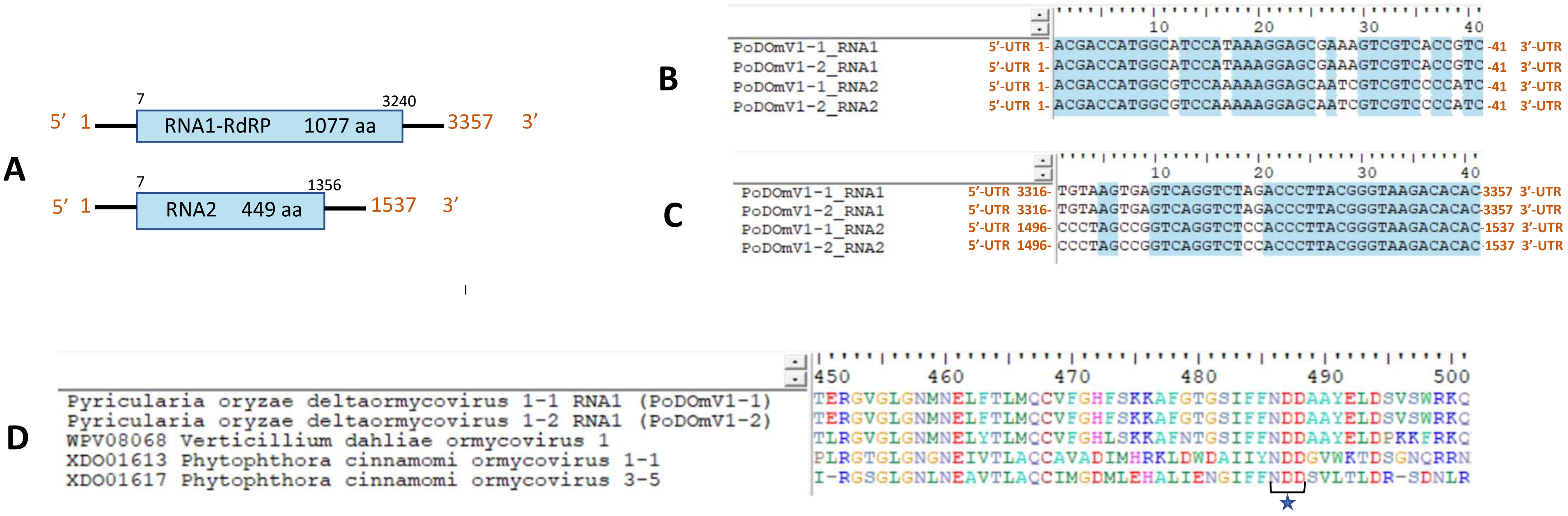

Supplementary Figure 2: Characterization of the Deltaormycovirus genome: (A) Genome organization of representative deltaormycoviruses, showing open reading frames (ORFs) and untranslated regions (UTRs) at the 5′ and 3′ termini, depicted as boxes and lines, respectively. (B–C) Sequence similarity between the 5′ (B) and 3′ (C) terminal regions of RNA1 and RNA2 segments from PoDOmV1-1 and PoDOmV1-2. (D) Multiple amino acid sequence alignment of RNA-dependent RNA polymerase (RdRp) from the identified viruses (PoDOmV1-1 and PoDOmV1-2) and representative members of the genus *Bormycovirus* available in GenBank, highlighting the conserved NDD motif at positions 486–488 within motif C

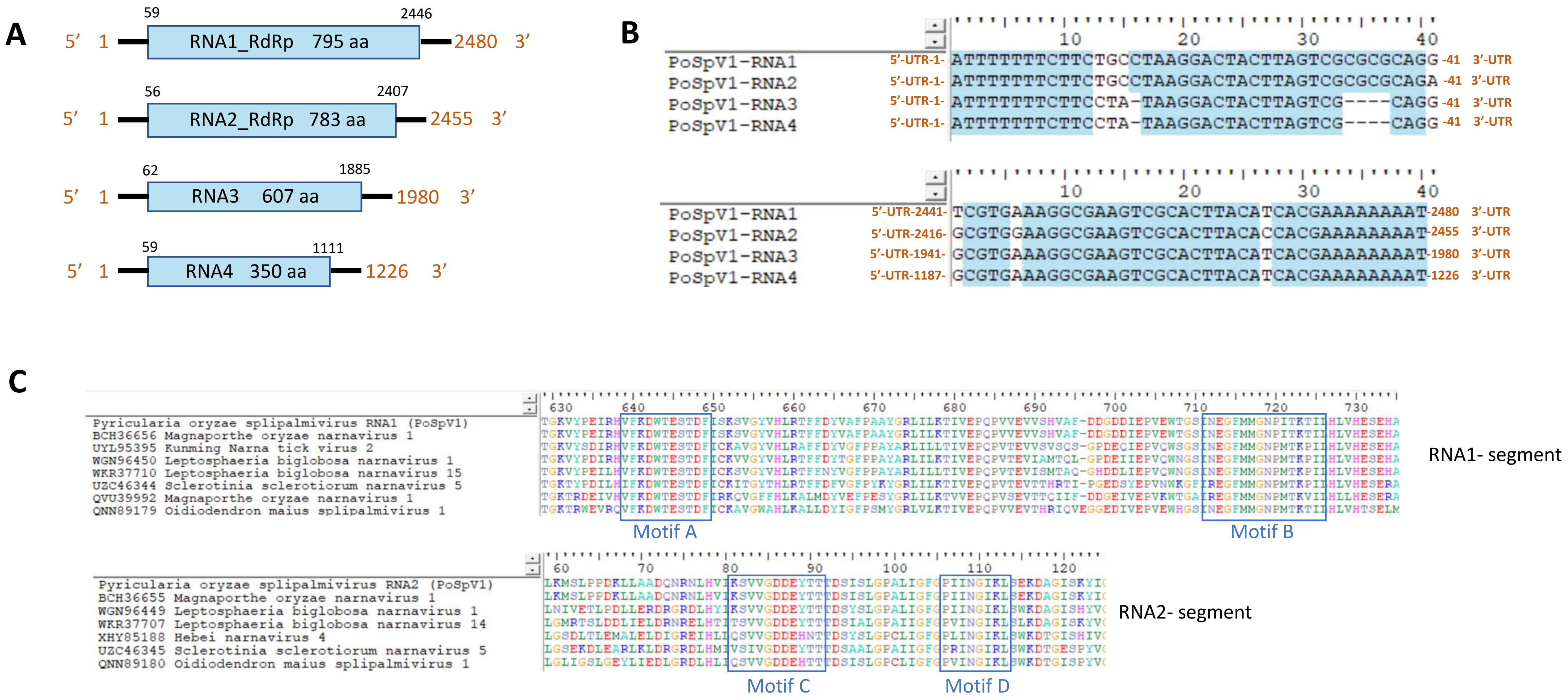


Supplementary Figure 3: Characterization of the Pyricularia oryzae splipalmivirus 1 (PoSpV1) genome: (A) Schematic representation of the PoSpV1 RNA genome, indicating open reading frames (ORFs) and untranslated regions (UTRs) at the 5′ and 3′ termini as boxes and lines, respectively. (B) Sequence similarity among the 5′ and 3′ terminal regions of the four genomic RNA segments. (C) Conserved RdRp motifs (A, B, C and D) distributed across the split RNA1 and RNA2 segments, highlighted in blue box.

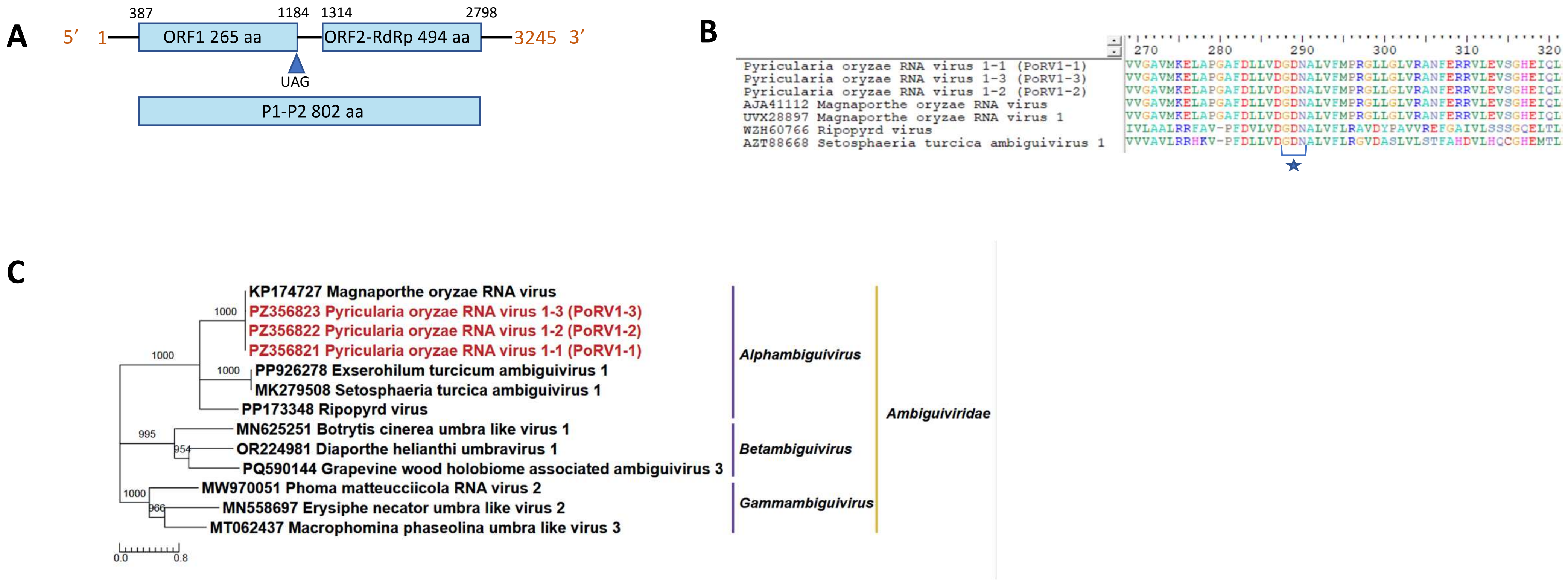


Supplementary Figure 4: Characterization of the PoRV1-1 genome: (A) Genome organization of PoRV1-1, with open reading frames (ORFs) and untranslated regions (UTRs) at the 5' and 3' termini represented as boxes and lines, respectively. ORF1 encodes a P1-P2 fusion protein (802 aa) generated via amber (UAG) stop codon readthrough. (B) Multiple amino acid sequence alignment of RdRps, highlighting the conserved GDN motif in motif C (blue star). (C) Maximum-likelihood phylogenetic analysis of predicted P1-P2 protein sequences from members of the *Ambiguiviridae* identified in this study (highlighted in red) and representative ICTV-recognized genomes from different genera available in GenBank.

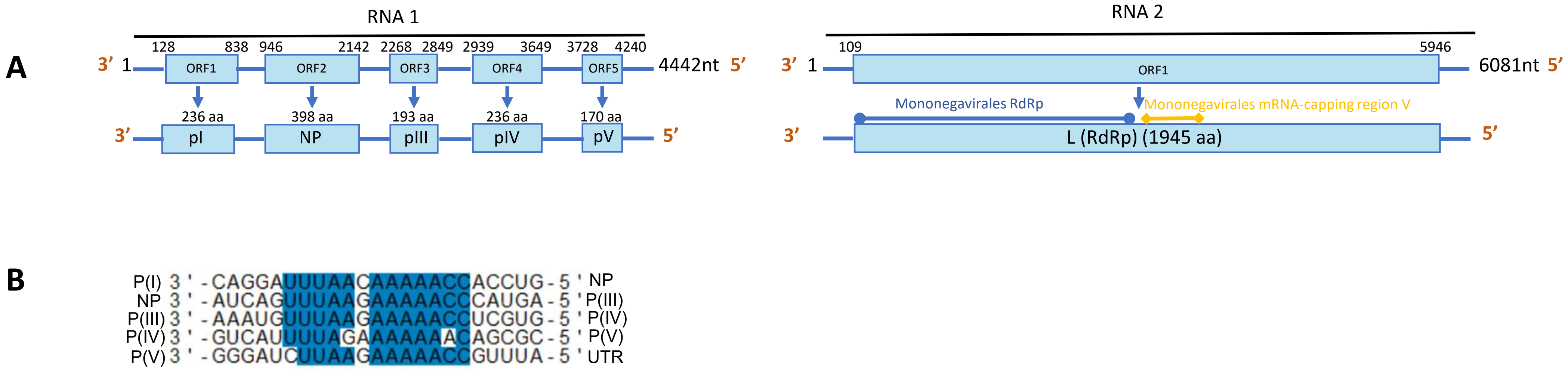


Supplementary Figure 5a: Characterization of the Pyricularia oryzae mymonavirus 1-1 (PoMV1-1) genome: (A) Schematic organization of the PoMV1-1 RNA genome, indicating the position and length of each ORF and their encoded proteins. Proteins are labelled with Roman numerals, except for the nucleoprotein (NP) and the L protein, which contains an RNA-dependant RNA polymerase domain (Accession pfam00946) and a conserved motif in region V required for mRNA capping (Accession pfam14318). (B) Alignment of putative gene-junction sequences between ORFs along the RNA1 segment in the 3'→ 5' orientation, with conserved motifs highlighted in blue.

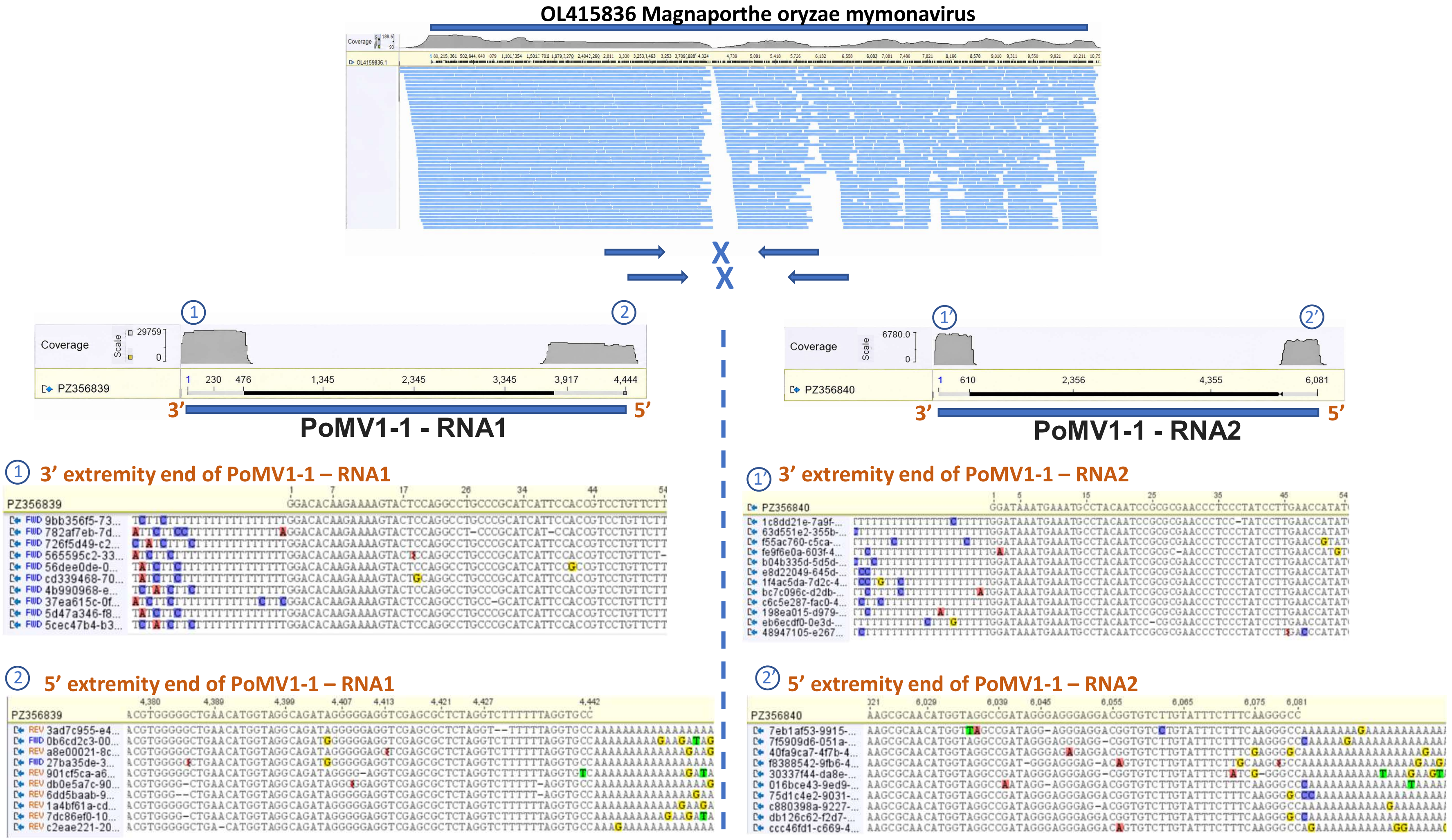


Representative subsamples of Nanopore reads from CH1184 and CH2061 mymonavirus mapped against the monopartite OL415836 genome (Magnaporthe oryzae mymonavirus), with the coverage along reference is shown for this subsamples.

PCR amplification using primers pairs flanking the putative segment jonction.

Nanopore read coverage obtained by RACE PCR and mapped to the 3' and 5' ends of both PoMV1-1 genomic segments.

Nanopore read alignement at the 3' end for both genomic segments.

Nanopore read alignement at the 5' ends for both genomic segments.

Supplementary Figure 5b: : Characterization of the PoMV1-1 genome and experimental evidence supporting its bi-segmented RNA genome.

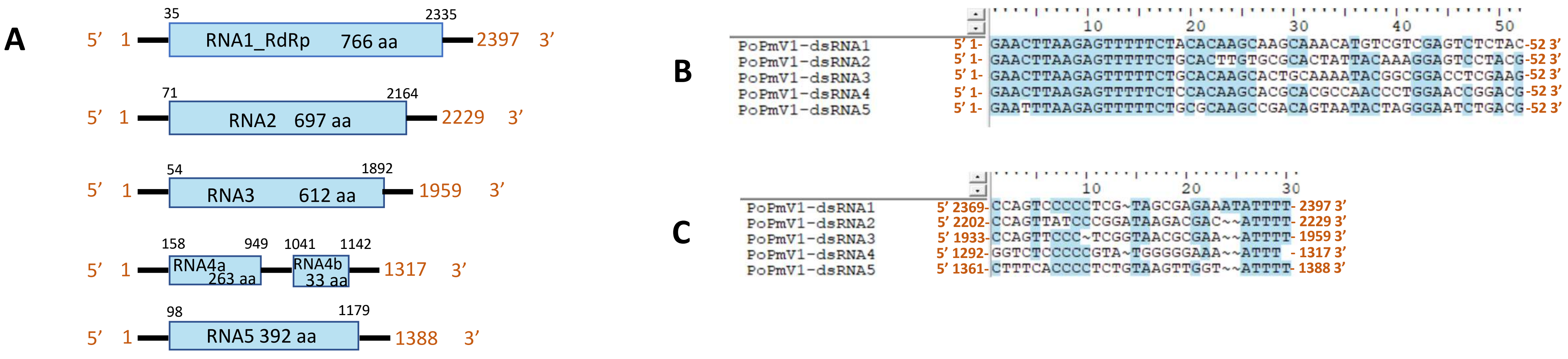


<u>Supplementary Figure 6</u>: Characterization of the PoPmV1 genome: (A) Schematic representation of the genome organization of PoPmV1, with open reading frames (ORFs) and untranslated regions (UTRs) depicted as open boxes and lines, respectively. (B-C) Sequence alignments of the 5′ (B) and 3′ (C) untranslated regions (UTRs) of dsRNA1-5 segments, highlighting conserved terminal features.
